# DnaJC7 specifically regulates tau seeding

**DOI:** 10.1101/2023.03.16.532880

**Authors:** Valerie A. Perez, David W. Sanders, Ayde Mendoza-Oliva, Barbara E. Stopschinski, Vishruth Mullapudi, Charles L White, Lukasz A. Joachimiak, Marc I. Diamond

## Abstract

Neurodegenerative tauopathies are caused by accumulation of toxic tau protein assemblies. This appears to involve template-based seeding events, whereby tau monomer changes conformation and is recruited to a growing aggregate. Several large families of chaperone proteins, including Hsp70s and J domain proteins (JDPs) cooperate to regulate the folding of intracellular proteins such as tau, but the factors that coordinate this activity are not well known. The JDP DnaJC7 binds tau and reduces its intracellular aggregation. However, it is unknown whether this is specific to DnaJC7 or if other JDPs might be similarly involved. We used proteomics within a cell model to determine that DnaJC7 co-purified with insoluble tau and colocalized with intracellular aggregates. We individually knocked out every possible JDP and tested the effect on intracellular aggregation and seeding. DnaJC7 knockout decreased aggregate clearance and increased intracellular tau seeding. This depended on the ability of the J domain (JD) of DnaJC7 to bind to Hsp70, as JD mutations that block binding to Hsp70 abrogated the protective activity. Disease-associated mutations in the JD and substrate binding site of DnaJC7 also abrogated its protective activity. DnaJC7 thus specifically regulates tau aggregation in cooperation with Hsp70.

## Introduction

Neurodegenerative tauopathies are caused by neuronal and glial accumulation of tau protein in amyloid fibrils^1^. We previously reported that tau has properties of a prion, in which assemblies of defined structure enter a cell, serve as a templates for their own replication, and lead to distinct patterns of neuropathology in a process termed “seeding”^2–5^. Recent cryo-electron microscopy studies have revealed distinct fibril morphologies (polymorphs) for several tauopathies, including Alzheimer’s Disease (AD), Progressive Supranuclear Palsy (PSP), and Corticobasal Degeneration (CBD)^6–8^. It has been known for decades that chaperones regulate intracellular protein aggregation. However, the factors that interact specifically with tau to regulate its assembly have not been comprehensively characterized.

Molecular chaperones regulate the folding, maturation, and degradation of proteins^9^. Heat shock proteins (HSPs) such as Hsp70 and Hsp90 have been reported to non-specifically regulate the folding of tau, alpha-synuclein, and other proteins implicated in neurodegenerative diseases^10^. A separate group of chaperones, the J-domain proteins (JDPs), bind myriad substrates through unclear mechanisms. JDPs shuttle substrates to Hsp70 and Hsp90^11,12^. They are defined by a highly conserved ∼70 amino acid J domain (JD) that enables binding to Hsp70 and stimulates its ATPase activity and subsequent transfer of the substrate to other regulatory components. JDPs thus play a key role in the Hsp70 protein folding/refolding cycle by linking substrates and chaperones.

JDPs have also been reported to directly modulate aggregation of tau and other neurodegenerative amyloid proteins^13,14^. We have previously determined the mode of tau:DnaJC7 interaction^15^. DnaJC7 binds tau with nanomolar affinity and sub-stoichiometrically reduces tau aggregation *in vitro*. Additionally, DnaJC7 preferentially binds inert tau, a form that does not spontaneously aggregate. We have now tested the specificity of tau binding to DnaJC7 vs. other JDPs and the role of Hsp70 in the regulation of intracellular tau aggregation.

## Results

### Identification of proteins that copurify with tau aggregates

Tau assemblies of distinct structure (“strains”) propagate indefinitely in clonal cultured cells that express the tau repeat domain (RD) containing two disease-associated mutations (P301L/V337M) fused to yellow fluorescent protein (YFP)^3,5^. To identify factors associated with tau, we studied the insoluble proteome of two clones, termed DS1 (which lacks inclusions) and DS10 (which propagates a unique strain). We first extracted DS1 and DS10 with sarkosyl to identify insoluble material. We boiled the insoluble fraction in SDS and resolved proteins by SDS-PAGE to isolate individual bands for extraction and analysis via mass spectrometry (Figure 1A, Supplementary Figure 1). We identified 12 unique proteins significantly enriched in DS10 compared to the DS1 control, and 49 proteins found only in the DS10 insoluble fraction (Figure 1B, C). These included VCP and Hsp70, which have previously been shown to modulate the tau aggregation process^16,17^, and other factors associated with protein quality control, autophagy, and the ubiquitin-proteasome system. As expected, the DS10 insoluble fraction was significantly enriched in tau and YFP. In contrast, the insoluble fraction of DS1 consisted predominantly of RNA-binding proteins and was de-enriched for tau and YFP.

**Figure 1.**
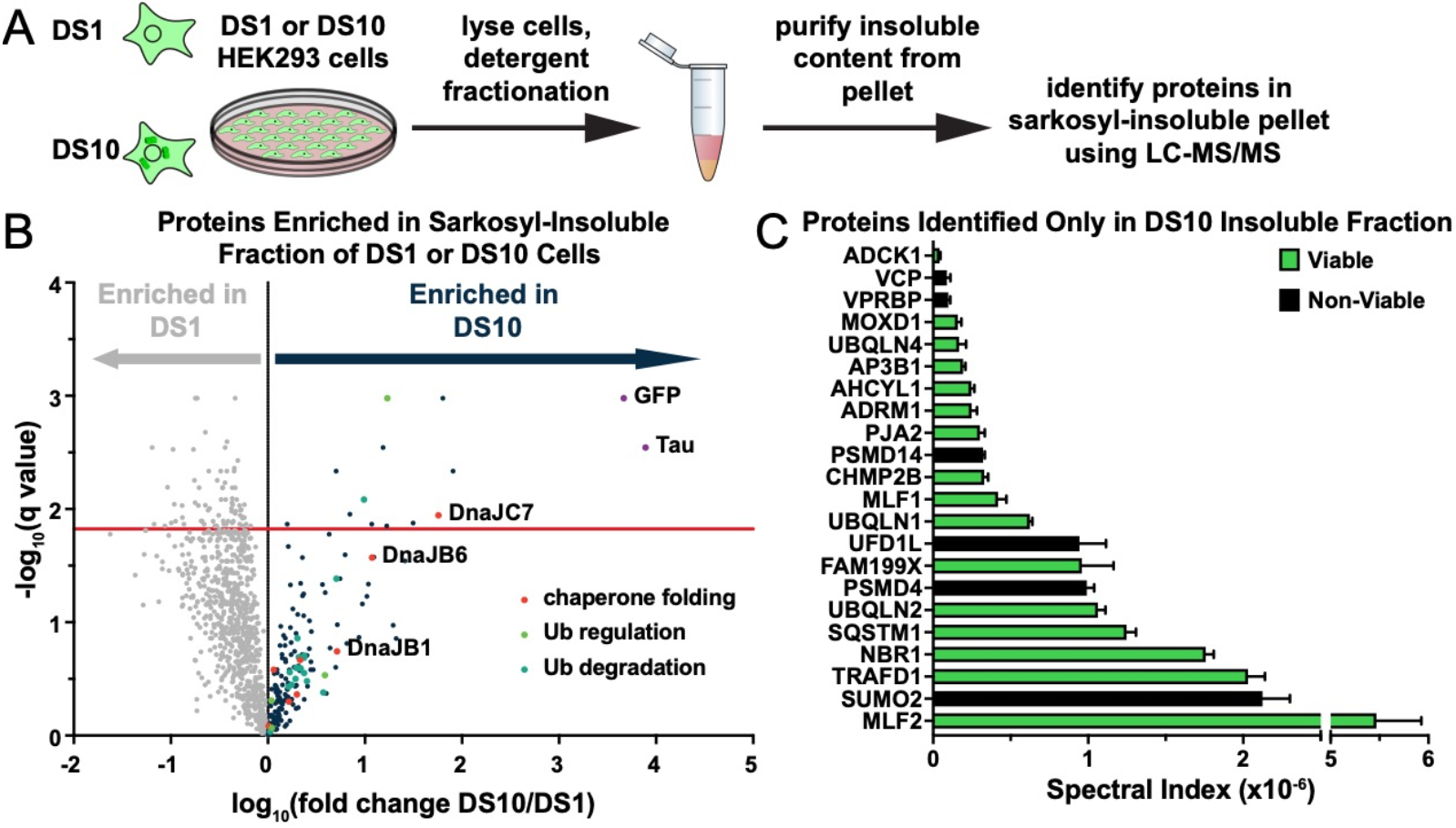
A proteomic approach to identify tau aggregate interactors. **A)** Tau aggresomes were partially purified from DS1 and DS10 HEK293 cells expressing tauRD-YFP. Detergent fractionation enabled generation of a sarkosyl-insoluble fraction containing tau aggresomes. Aggregates were recovered from the pellet by running protein on an SDS-PAGE gel and then extracting protein from individual lanes from the gel to be analyzed via LC-MS/MS. **B)** Volcano plot showing proteins enriched in the sarkosyl-insoluble fraction as a fold enrichment from cells expressing tauRD-YFP aggregates (DS10, dark blue dots) over cells expressing tauRD-YFP that does not form aggregates (DS1, gray dots). The red line indicates an FDR of 1.5%. GO term enrichment analyses of biological processes is also shown for select GO terms: orange dots, chaperone-mediated protein folding (chaperone folding); green dots, regulation of ubiquitination (Ub regulation); teal dots, ubiquitin-dependent protein catabolic process (Ub degradation). **C)** Spectral indices for a selection of the proteins identified only in the DS10 insoluble fraction. Viable knockouts are shown as green bars. Non-Viable knockouts are shown as black bars. Error bars represent the SEM of three extracted protein SDS-PAGE gel bands.

**Figure 1 – Supplement 1.**
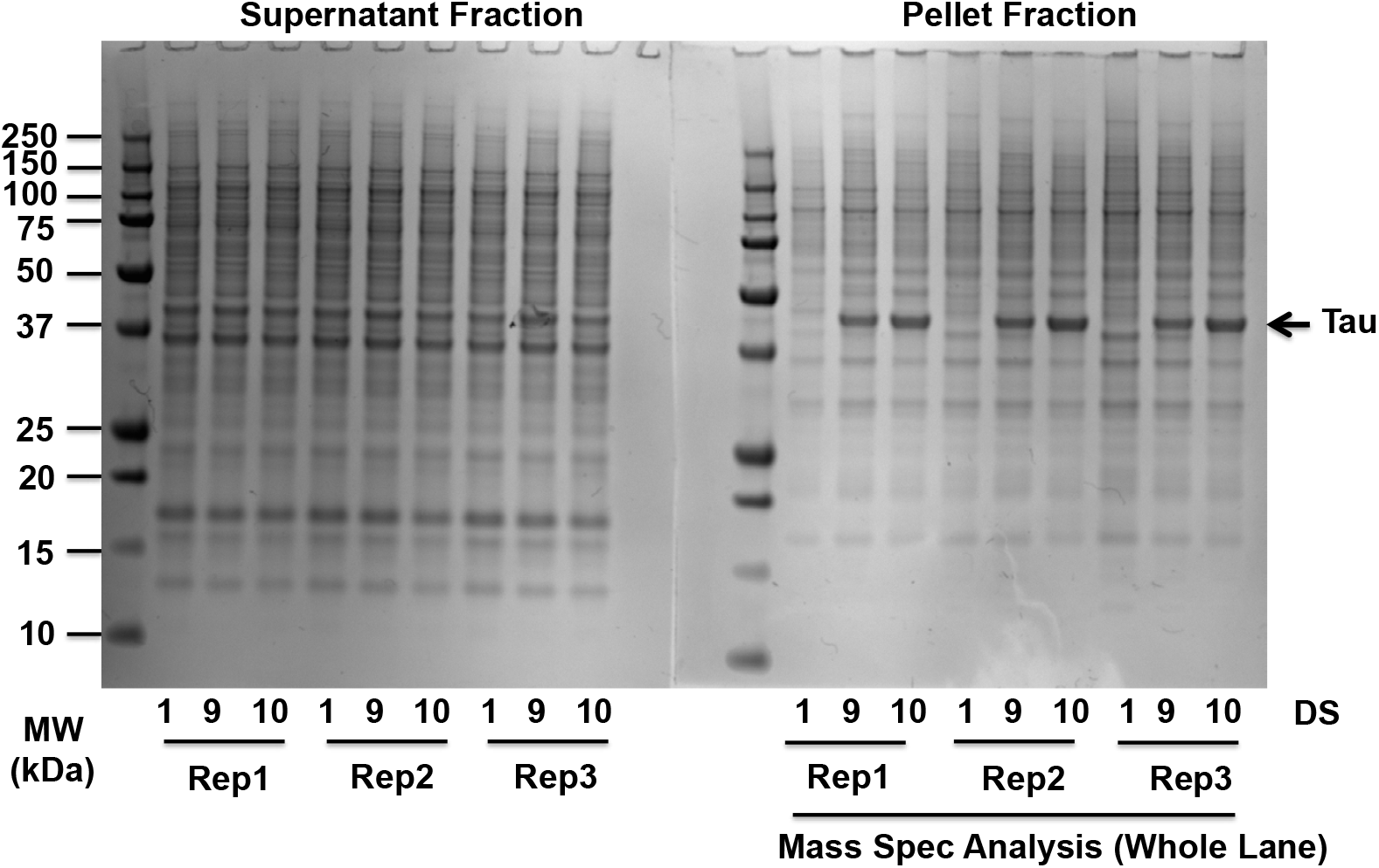
Partial purification of tau aggresomes. Sarkosyl-soluble and sarkosyl-insoluble fractions were run on gels and proteins were stained with SimplyBlue protein stain. DS10 (contains tau aggregates) but not DS1 (lacks tau aggregates) insoluble fractions featured a significant enrichment of tau RD-YFP. Whole lanes for pellet fractions were analyzed by mass spectrometry (biological triplicates). The DS9 strain cell line is included as a tau-aggregate containing control. Source data for this figure are provided in Figure 1 - Supplement 1 - Source Data 1.

### DnaJC7 knockout reduces clearance and increases inclusion density

We first determined whether any of the top interactors from the proteomic screen would modulate tau aggregate clearance. We utilized a cell line that propagates the DS10 strain with tauRD-YFP expression regulated by a tetracycline-repressible (tet-off) promoter. These cells (henceforth termed OFF1::DS10) constitutively produce large juxtanuclear tau aggregates, whose clearance can be monitored by loss of YFP fluorescent puncta after addition of doxycycline to the cell media to shut off gene expression (Figure 2A).

**Figure 2.**
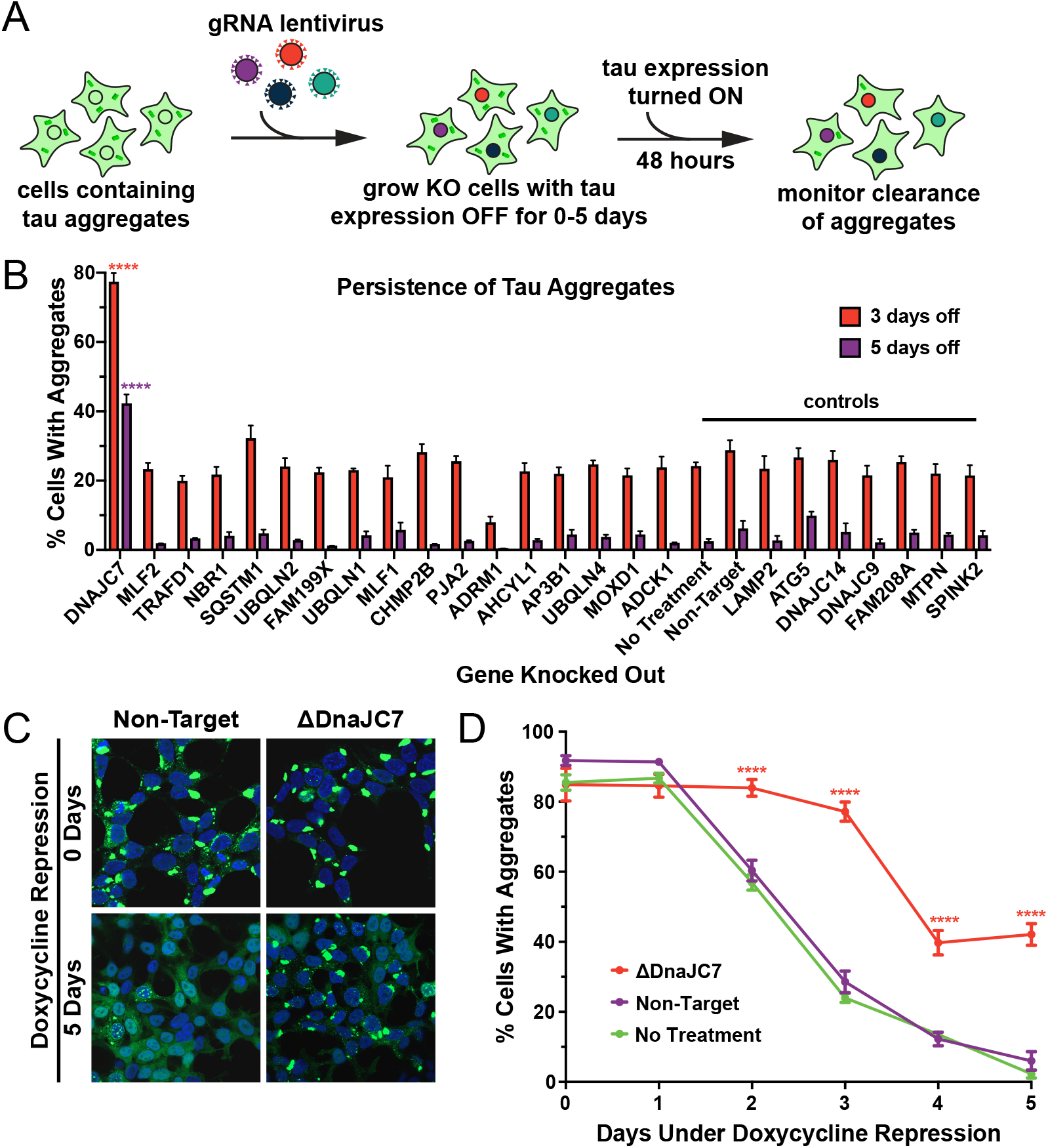
DnaJC7 KO uniquely extends tau seed lifespan in dividing cells. **A)** Schematic showing the HEK293 OFF1::DS10 system. A selection of the hits from the proteomics screen were knocked out in these cells. The cells are then allowed to grow with tau expression turned OFF for 0-5 days before resuming tau expression. Error bars represent the SEM of six technical replicates. **B)** The persistence of tau aggregates in OFF1::DS10 cells with the indicated knockout was quantified following 3 (orange bars) or 5 (purple bars) days of repressed tau expression. Error bars represent the SEM of six technical replicates. **C)** Confocal microscopy images showing tau aggregate organization in the DnaJC7 KO and nontargeting control cells following 0 or 5 days of repression of tau expression. **D)** Extended time course for tau aggregate clearance in the OFF1::DS10 system with DnaJC7 KO (orange) and the nontargeting (purple) and untreated (green) controls. Error bars represent SEM of six technical replicates. ^*^ = p<0.05, ^**^ = p<0.01, ^***^ = p<0.001, ^****^ = p<0.0001

**Figure 2 – Supplement 1.**
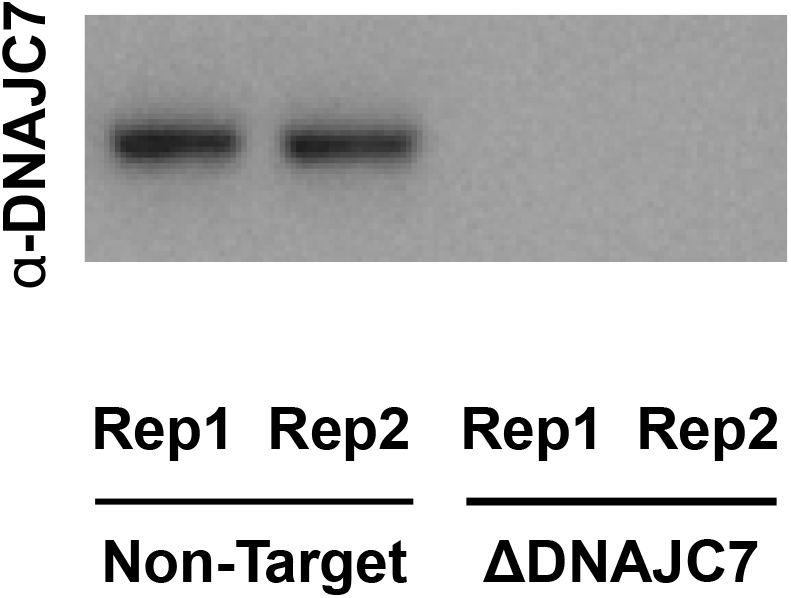
Western blot confirms DnaJC7 KO in (OFF1::DS10) cells. Immunoblotting for DnaJC7 confirms that DnaJC7 is knocked out in the Tet-regulated tauRD-YFP aggregate line (OFF1::DS10). Source data for this figure are provided in Figure 2 - Supplement 1 - Source Data 1.

We knocked out a selection of genes enriched in or found only in the DS10 insoluble fraction plus control genes in the OFF1::DS10 cell lines and monitored tau clearance. Knockout of certain genes (e.g., VCP, SUMO2, black bars in Figure 1C) was lethal, and thus their effects could not be tested. We shut off tau expression for three or five days to identify cells which had fully cleared aggregates and manually counted cells to quantify the effects of the knockouts on tau aggregate clearance. OFF1::DS10 cells with DnaJC7 KO out had the greatest number of cells still containing tau aggregates after three days (∼80% containing aggregates) or five days (∼40% containing aggregates) of repression (Figure 2B). This contrasted sharply to the rest of the knockout cell lines generated, which all cleared most of the aggregates after five days of repression. Additionally, confocal microscopy revealed that DnaJC7 KO increased inclusion density and decreased the quantity of extra-aggresomal (diffuse) tau compared to the nontargeting control (Figure 2C). DnaJC7 knockout was confirmed by Western Blot (Supplementary Figure 2).

To test for DnaJC7-mediated clearance of invisible seeds, we shut off tau expression for zero to five days and then restarted expression for two days. DnaJC7 KO decreased the rate of tau aggregate clearance, with ∼40% of cells contained aggregates after five days of repression vs. ∼0% of the nontargeting and untreated control cells (Figure 2D). The profound effects of DnaJC7 indicated that it likely played a key role in mediating seed clearance.

### DnaJC7 and DnaJB6 uniquely regulate tau aggregation

Given that DnaJC7 KO impeded the clearance of tau aggregates, we next tested whether this would modify seeded tau aggregation. We have previously developed a biosensor cell line that stably expresses tauRD (P301S) linked to the mClover3 or mCerulean3 fluorescent proteins^18,19^. Application of exogenous aggregates induces intracellular tau aggregation that is quantified by fluorescence resonance energy transfer (FRET). Exogenous fibrils bind to heparan sulfate proteoglycans (HSPGs) on the cell surface and trigger their own uptake via macropinocytosis. Internalized tau seeds then escape endolysosomal trafficking, enter the cytoplasm, and trigger further intracellular aggregation^20^. Alternatively, aggregates can be introduced directly into the cytoplasm by cationic lipids such as Lipofectamine 2000, which increases the induced seeding efficiency.

To test the specificity of DnaJC7, we used CRISPR/Cas9 to individually knock out all known JDPs in the biosensor line. We used gRNAs from the Brunello library^21^ in pools of four for each gene, to produce 50 unique JDP knockout biosensor cell lines (Figure 3A). We exposed the knockout biosensors to naked seeds at various concentrations. After 48 h we analyzed seeding in the cells via flow cytometry. Only DnaJC7 and DnaJB6 KO significantly increased tau seeding relative to the nontargeting control (Figure 3B, full dose titrations shown in Supplementary Figure 3A).

**Figure 3.**
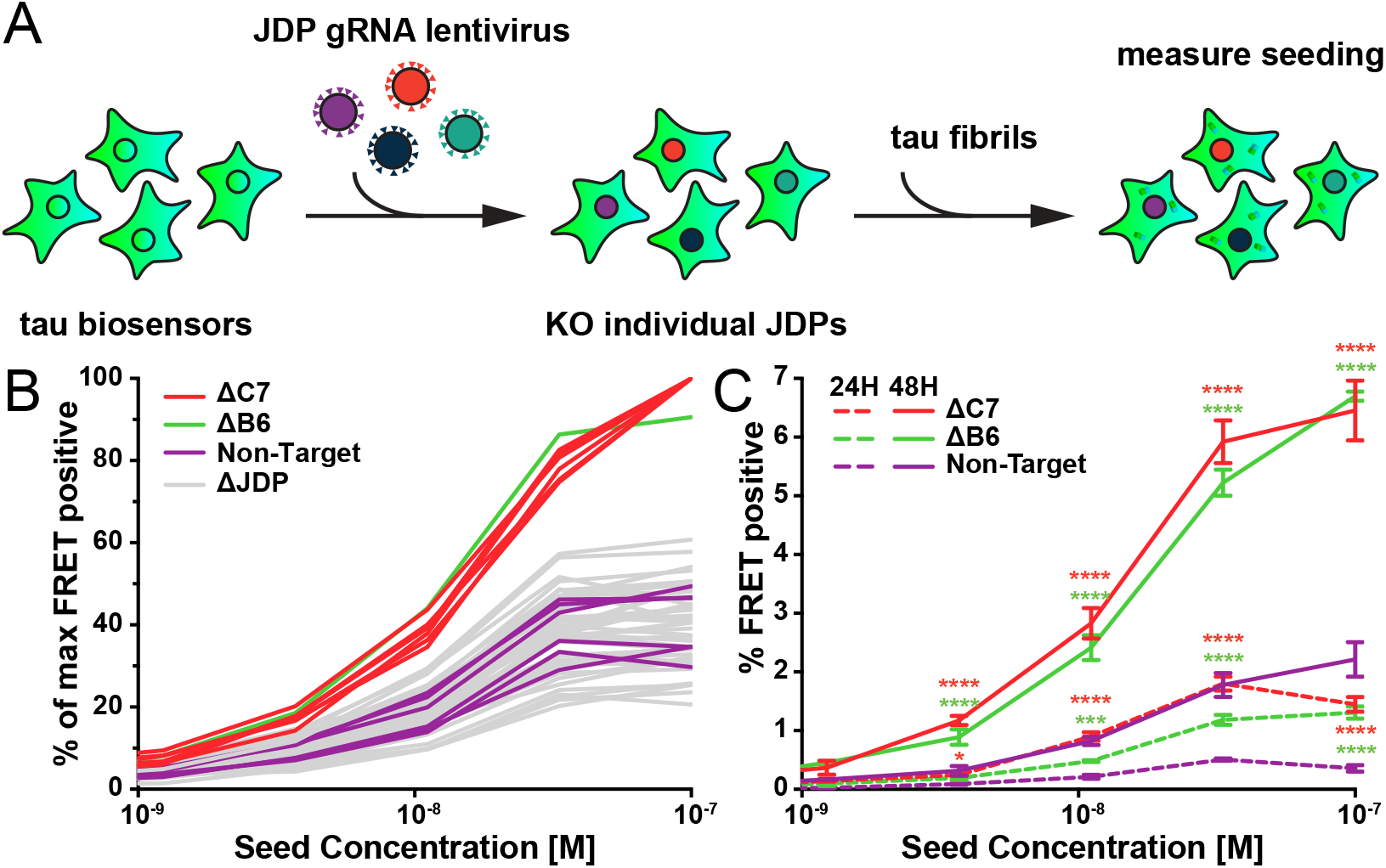
A targeted genetic screen for modifiers of transient tau seeding identifies specific JDPs. **A)** Schematic showing the HEK293 tau biosensor system consisting of tauRD fused to either mCerulean3 or mClover3 fluorescent proteins. The base biosensor cells had each JDP individually knocked out to generate 50 distinct cell lines. Recombinant, sonicated tau fibrils (seeds) are added to the cells to induce seeding of the tauRD constructs, which is detected as a FRET signal via flow cytometry 48 hours after treatment. **B)** Representative data showing the effects of the individual knockouts of JDPs on tau seeding in biosensor cells, quantified as FRET signal via flow cytometry. Cells were seeded with a dose titration of sonicated tau fibrils. Knockout of DnaJC7 (ΔC7, orange) and DnaJB6 (ΔB6, green) are highlighted. The remaining JDP knockouts are denoted as ΔJDP (gray). Each batch of knockout cell lines was normalized to the DnaJC7 KO seeding signal and then all batches are plotted together. The seeding assay for the DnaJC7 KO and the nontargeting control (Non-Target, purple) were repeated ten times. **C)** Extended time course harvesting of the tau seeding assay for DnaJB6 KO, DnaJC7 KO, and nontargeting control cells harvested at 24 h (24H, dashed lines) and 48 h (48H, solid lines) timepoints. Coloring as in **B)**. Error bars represent SEM of three technical replicates. ^*^ = p<0.05, ^**^ = p<0.01, ^***^ = p<0.001, ^****^ = p<0.0001

**Figure 3 – Supplement 1.**
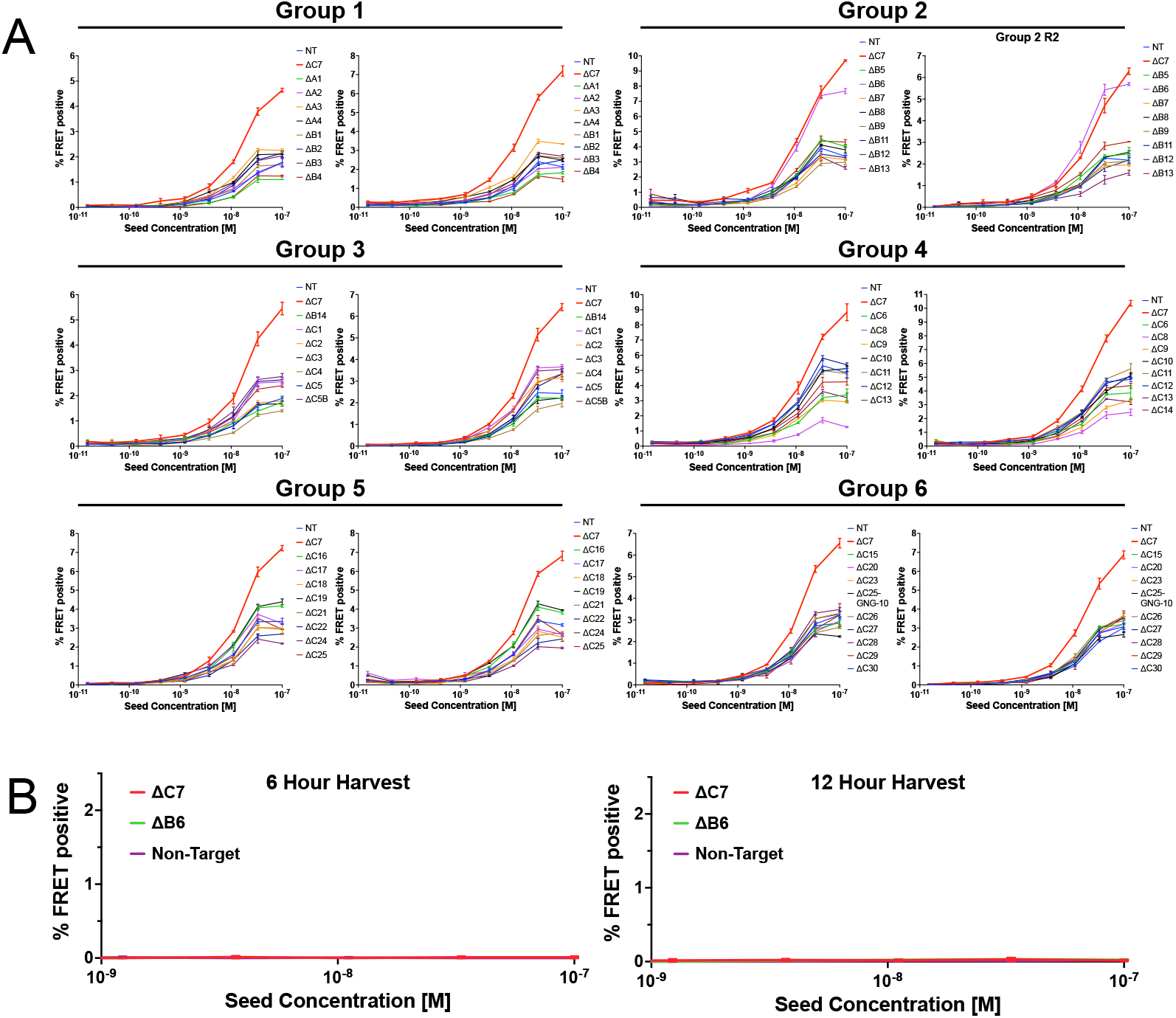
All individual groups of the JDP CRISPR screen. **A)** Full tau dose titrations for all batches of individual knockouts of JDPs on tau seeding in biosensor cells, quantified as FRET signal via flow cytometry. Cells were seeded with a dose titration of sonicated tau fibrils. Knockout of DnaJC7 (ΔC7, red) and the nontargeting control (NT, blue) are highlighted in each batch. **B)** Extended time course harvesting of the tau seeding assay for DnaJC7 KO (ΔC7, orange), DnaJB6 KO (ΔB6, green), and nontargeting control (Non-Target, purple) cells harvested at 6 h and 12 h timepoints. All error bars represent SEM of three technical replicates.

We also tested whether DnaJB6 and DnaJC7 KO changed the kinetics of intracellular seeding by evaluating seeding at 6, 12, 24, and 48 h timepoints. We observed no significant seeding in any cell lines at 6 and 12 h (Supplementary Figure 3B). At 24 h, DnaJC7 KO enabled more intracellular aggregation than DnaJB6 KO. Both knockout cell lines exhibited higher seeding than the nontargeting control cell line. At 48 h DnaJC7 and DnaJB6 KO cell lines had comparable seeding ∼3-fold higher than the nontargeting control.

### DnaJC7 KO increases seeding from tauopathy brains

To further characterize DnaJC7 and DnaJB6, we tested their effect on brain-derived tau seeding. We treated DnaJC7 and DnaJB6 KO cell lines with either recombinant tau fibrils or brain lysates from subjects with Alzheimer’s Disease (AD), Progressive Supranuclear Palsy (PSP), or Corticobasal Degeneration (CBD). We exposed biosensors directly to 25 μL of brain lysates or transfected 5 μL of brain lysates using Lipofectamine 2000. Additionally, biosensors were treated with 20 μg total protein of DS cell lysates or transfected with 5 μg of cell lysates using Lipofectamine 2000. Only DnaJC7 KO significantly increased seeding of the homogenates (Figure 4A, B). DnaJC7 KO alone enhanced seeding from naked AD lysate, whereas control and DnaJB6 KO cell lines did not. Thus, DnaJC7 was not constrained by the specific conformation of tau seeds.

**Figure 4.**
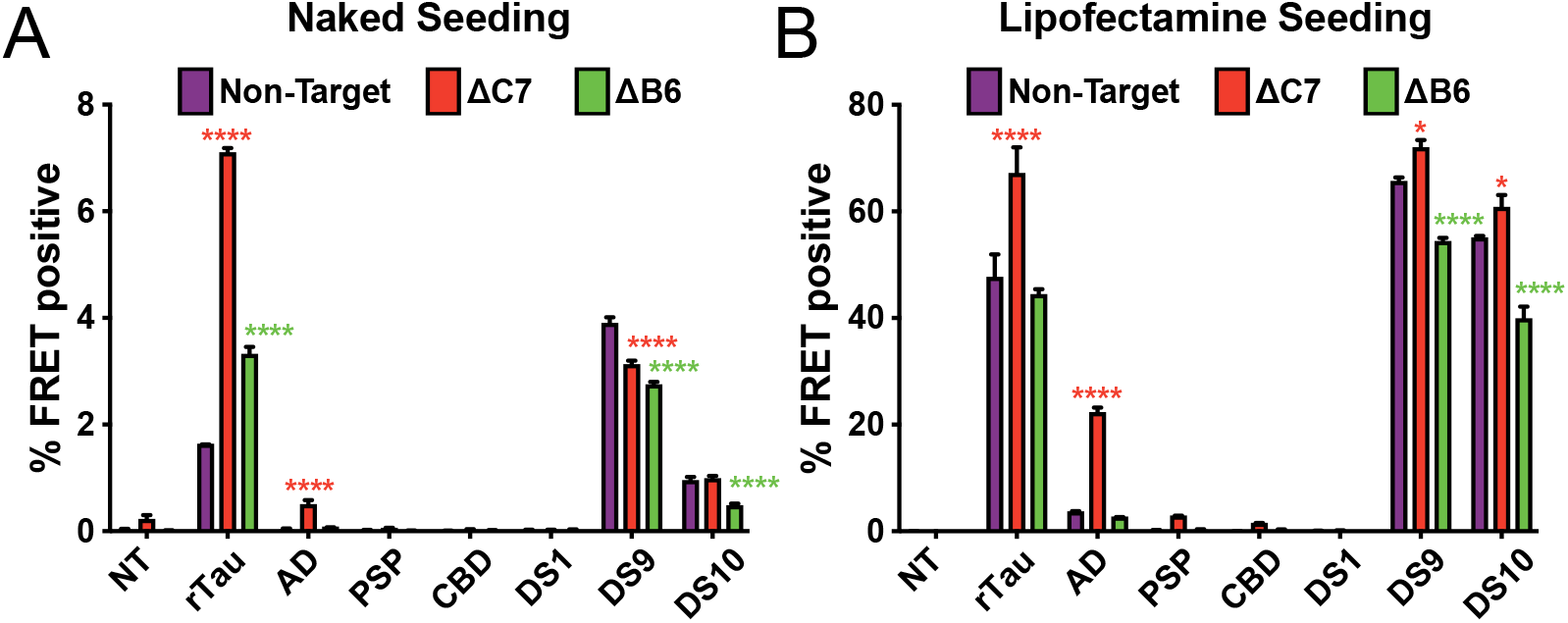
DnaJC7 knockout increases tau seeding across multiple seed sources. Intracellular **A)** naked or **B)** lipofectamine-mediated tau seeding in nontargeting control (Non-Target, purple), DnaJC7 KO (ΔC7, orange), and DnaJB6 KO (ΔB6, green) tau biosensor cells. Cells were seeded with sonicated recombinant tau fibrils (rTau); brain lysates from patients with Alzheimer’s Disease (AD), Progressive Supranuclear Palsy (PSP), or Corticobasal Degeneration (CBD); and cell lysates of DS1, DS9, or DS10 cells. 100 nM or 10 nM of recombinant tau were added for naked and lipo seeding, respectively. 25 μL or 5 μL of patient brain or lysate or 20 μg or 5 μg cell lysate were added for naked and lipo seeding, respectively. NT denotes no treatment with seeds. Error bars represent SEM of three technical replicates. ^*^ = p<0.05, ^**^ = p<0.01, ^***^ = p<0.001, ^****^ = p<0.0001

### DnaJC7 overexpression rescues knockout tau biosensors

To test for specificity of knockout of DnaJC7 in the biosensor cells, we transiently overexpressed DnaJC7 using a Ruby-DnaJC7 fusion with a coding sequence resistant to targeting by the gRNAs used to generate the original DnaJC7 KO cell line (Supplementary Figure 4A). Expression of the Ruby-DnaJC7 fusion constructs was confirmed by Western Blot (Supplementary Figure 4B). Ruby-DnaJC7 overexpression in the DnaJC7 KO cells reduced tau seeding relative to Ruby alone and vehicle controls (Figure 5A). Overexpression of Ruby-DnaJC7 in the control tau biosensors without DnaJC7 KO modestly reduced tau seeding (Figure 5B).

**Figure 5.**
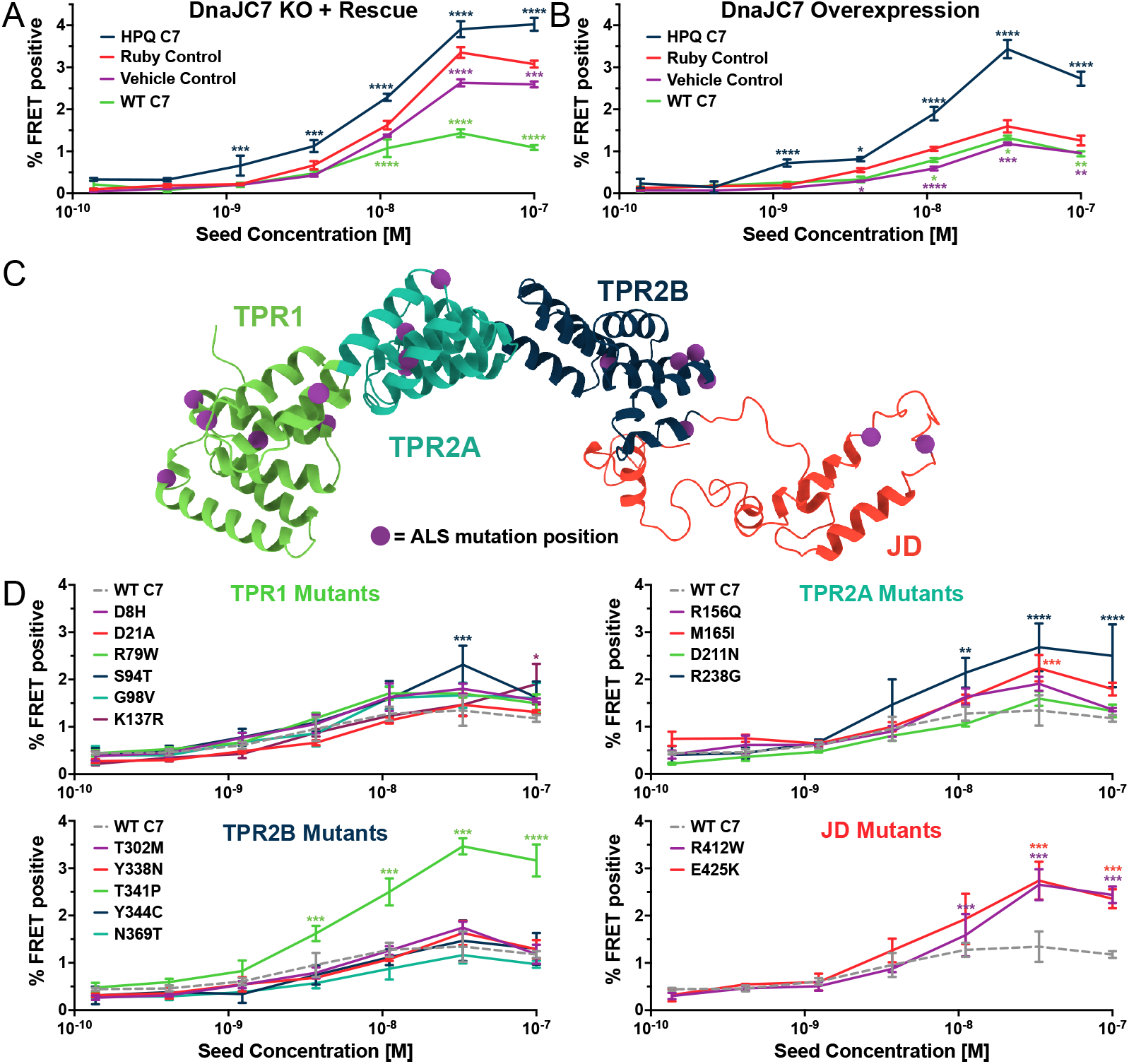
DnaJC7 regulates tau seeding in multiple experimental approaches. **A)** Rescue of DnaJC7 KO in tau biosensor cells with either wildtype (WT C7, green) or HPQ mutant (HPQ C7, dark blue) DnaJC7 constructs. The Ruby fluorophore alone (Ruby Control, orange) and a vehicle control (Vehicle Control, purple) were also added to the DnaJC7 KO cells. **B)** Overexpression of DnaJC7 constructs in control tau biosensor cells. The cells were transfected with the same constructs as in **A). C)** Model of DnaJC7 with domains colored as follows: TPR1, green; TPR2A, teal; TPR2B, dark blue; JD, orange. Positions of ALS-associated mutations are shown as purple spheres. **D)** Rescue of DnaJC7 KO in tau biosensor cells with ALS-associated mutants of DnaJC7 and WT control, sorted by domain location. Rescue with the WT DnaJC7 construct is shown in all domains as a grey, dashed line. Error bars represent SEM of three technical replicates. ^*^ = p<0.05, ^**^ = p<0.01, ^***^ = p<0.001, ^****^ = p<0.0001

**Figure 5 – Supplement 1.**
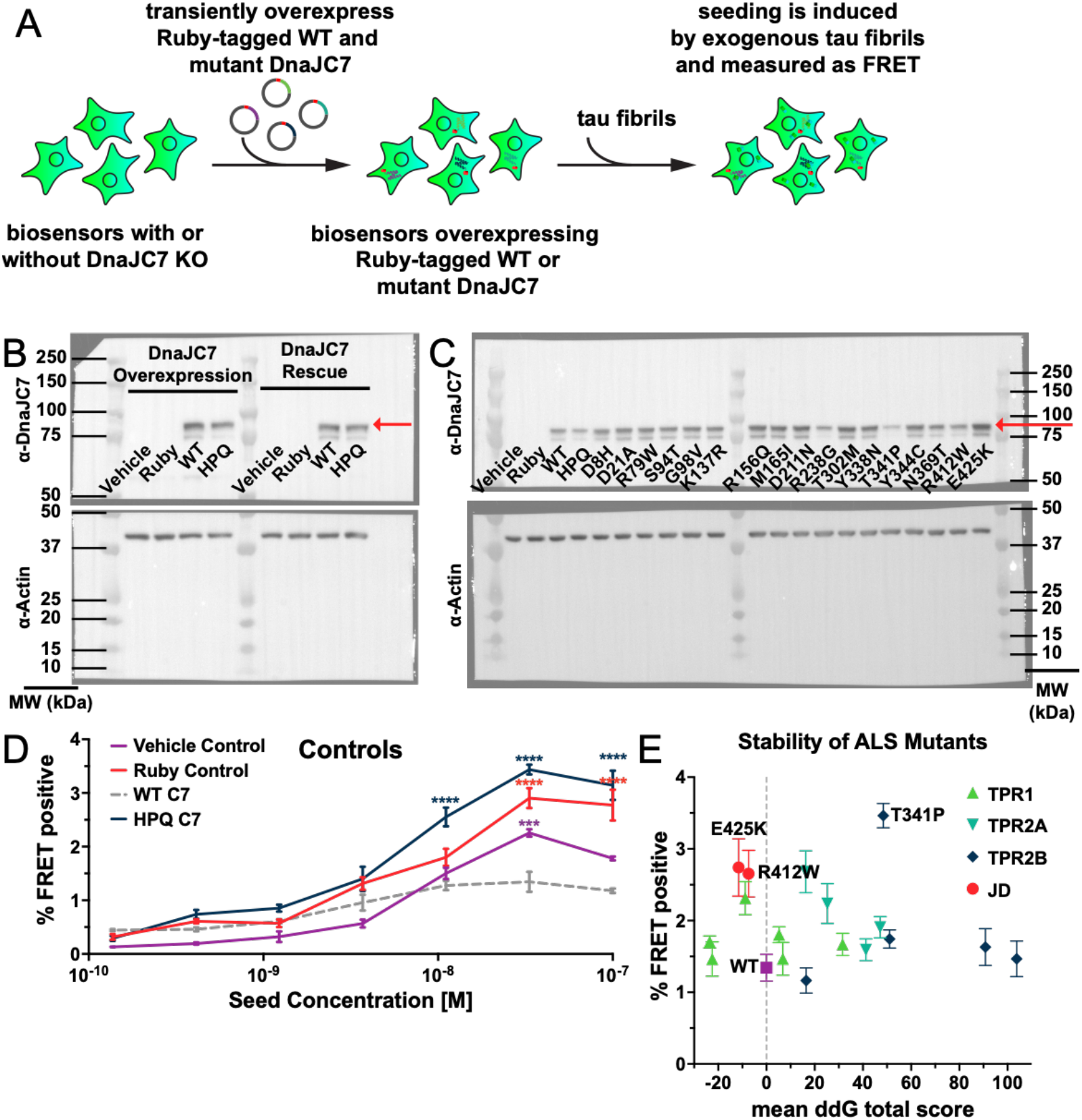
Stability and expression of DnaJC7 mutants in tau biosensors. **A)** Schematic showing tau biosensor cells with or without DnaJC7 KO transiently overexpressing different Ruby-tagged gRNA-resistant DnaJC7 constructs. Cells are allowed to express the constructs for two days before being plated for the seeding assay. **B)** Immunoblotting for DnaJC7 confirms expression of the Ruby-WT DnaJC7 and Ruby-DnaJC7 (HPQ) mutant constructs in tau biosensor cells without (Overexpression) and with endogenous DnaJC7 knocked out (Rescue). Ruby fusion constructs are highlighted by a red arrow. **C)** Immunoblotting for DnaJC7 confirms expression of the Ruby-WT DnaJC7 and Ruby-DnaJC7 ALS mutant constructs in tau biosensor cells with endogenous DnaJC7 knocked out. Ruby fusion constructs are highlighted by a red arrow. **D)** Positive and negative controls utilized in the rescue of DnaJC7 KO in tau biosensor cells with ALS-associated mutants of DnaJC7, colored as follows: Vehicle control, purple; Ruby control, orange; WT DnaJC7, grey dashed; HPQ mutant, dark blue. Error bars represent SEM of three technical replicates. **E)** Rosetta-calculated mean Gibbs free energy shift (ddG) of the ALS-associated mutants of DnaJC7 vs their rescue seeding with 33 nM of tau fibrils. Grey dashed line denotes a mean ddG total score of 0. Mutants are colored according to their domain localization: TPR1, green; TPR2A, teal; TPR2B, dark blue; JD, orange. Error bars represent SEM of three technical seeding replicates. ^*^ = p<0.05, ^**^ = p<0.01, ^***^ = p<0.001, ^****^ = p<0.0001. Source data for this figure are provided in Figure 5 - Supplement 1 - Source Data 1.

To test the role of DnaJC7 binding to Hsp70, we introduced a point mutation into the Ruby-DnaJC7 fusion that inhibits Hsp70 binding and substrate handoff^22,23^. The aspartic acid in the histidine-proline-aspartic acid (HPD) motif was mutated to a glutamine (D411Q, henceforth termed HPQ mutant) to prevent the Hsp70 interaction. Expression of Ruby-DnaJC7 (HPQ) in the DnaJC7 KO biosensors did not rescue the WT seeding phenotype and instead enhanced tau seeding (Figure 5A). Surprisingly, overexpression of DnaJC7 (HPQ) in the scrambled control tau biosensors also increased tau seeding, consistent with a dominant negative effect (Figure 5B).

### Disease-associated DnaJC7 mutations differentially modulate tau seeding

DnaJC7 mutations cause dominantly inherited Amyotrophic Lateral Sclerosis (ALS)^24^. 17 mutations have been identified that are distributed across all domains of DnaJC7 (Figure 5C). The TPR1 and TPR2A domains have been implicated in binding to EEVD motifs in Hsp70 and Hsp90, respectively^25^. We have previously found that the TPR2B domain mediates DnaJC7 binding to tau^15^. Additionally, DnaJC7 was recently shown to bind the prion-like domain of the ALS-associated protein TDP-43 and mitigate its ability to phase separate in vitro^26^. To test whether these disease-associated mutations can also affect how DnaJC7 modulates tau seeding, we introduced each mutation individually into the Ruby-DnaJC7 construct. We then transiently transfected them into the DnaJC7 KO tau biosensor cell line and quantified changes in tau seeding capacity (Supplementary Figure 4A).

Rescue of DnaJC7 KO with most of the mutants suppressed seeding similar to WT. Five mutants increased tau seeding vs. WT (Figure 5C). The five inhibitory mutations were in all four domains of DnaJC7, with two located in the J domain (R412W and R425K). Neither J domain mutant resembled the seeding phenotype of DnaJC7 (HPQ). However, the T341P mutant recapitulated the effect of the DnaJC7 (HPQ) mutant, suggesting a toxic gain of function with effects similar to blocking DnaJC7:Hsp70 interactions. We ruled out mutant effects on DnaJC7 expression via Western Blot (Supplementary Figure 4C). The expression of most of the constructs resembled WT, while those that increased tau seeding were slightly reduced in expression, consistent with a gain-of-function effect. Thus, DnaJC7 may function within a more complex network of chaperones to control tau aggregation.

## Discussion

This study began with an unbiased proteomic screen to identify factors that co-purify with insoluble tau from cells stably propagating a tau strain (DS10). We confirmed the top hits and identified DnaJC7 as a unique regulator of tau aggregation and clearance. The large family of JDPs are thought to be functionally redundant. However, when we tested the specificity of all members of the JPD family using a candidate CRISPR/Cas9 genetic knockdown approach, we identified DnaJC7 and to a lesser degree DnaJB6 as unique regulators of tau aggregation. Finally, we tested newly identified mutations in DnaJC7 which cause ALS and found that mutations in the J domain have dominant negative effects on tau aggregation. This is consistent with the idea that Hsp70-based coordination of DnaJC7 is central to its activity and is linked to protein aggregation in ALS (which usually doesn’t feature tauopathy) and to regulation of tau.

### Disease-associated mutations of DnaJC7 differentially affect tau seeding

Pathogenic mutations in several JDPs have previously been linked to multiple heritable neurodegenerative diseases as well as diseases targeting other organ systems^27^. These mutations, ranging from missense to stop-gain mutants, inhibit the ability of the JDPs to bind clients or transfer them to Hsp70 and produce different disease pathologies. Recently, a series of protein truncating variants and missense mutations to the DnaJC7 gene have been identified as causal for ALS^24^. The missense mutations were found across all domains of DnaJC7, with most localized to the helix-turn-helix loops on the TPR motifs.

DnaJC7 rescue experiments to test 17 known ALS-associated missense mutations revealed that the T341P mutant recapitulated the seeding profile of the DnaJC7 (HPQ) mutant incapable of binding to Hsp70. The T341P mutation is in the TPR2B domain, which we have previously found to be the main site of tau binding on DnaJC7^15^. Additionally, rescue with the two J domain mutants (R412W and R425K) increased seeding relative to the WT DnaJC7 sequence, but both mutants still seeded lower than the T341P and HPQ mutants. Further, the S94T mutant, which may impede Hsp90 binding, also moderately increased tau seeding relative to WT. ALS-associated mutations may inhibit DnaJC7 interaction with other chaperones (i.e., Hsp70 and Hsp90) and thus impair substrate (tau) handoff into the chaperone-mediated protein refolding cycle. This agrees with previous observations that DnaJC7 cooperates with other chaperones to mitigate FUS and TDP-43 toxicity in yeast and *in vitro*^28,26^.

To further test the perturbations to the DnaJC7 structure afforded by the ALS-associated mutations, we used Rosetta to calculate the predicted change in structural stability of the monomer in response to each mutation^29^. We found no significant correlation between the predicted energy perturbation of the mutations and the resulting seeding phenotype (Supplemental Figure 4E). Although the TPR2B mutations Y338N and Y344C are predicted to have the highest destabilizing effects on DnaJC7, they resembled WT in regulating seeding. In contrast, the two J domain mutants were predicted to stabilize the DnaJC7 structure but increased seeding. Instead of destabilizing DnaJC7, these mutations exhibit gain-of-function effects likely through dysregulation of interactions within the chaperone network. Disease-associated mutations occur in both DnaJC7 and DnaJB6^24,30,31^. This suggests that DnaJB6, DnaJC7 and other JDPs are not functionally redundant.

### DnaJC7 cooperates with Hsp70 to regulate intracellular tau aggregation

Chaperones function in networks to regulate the folding of a myriad of substrates. The Hsp70 chaperones need JDPs to provide client specificity and activate their ATPase activity. Additionally, co-chaperones such as the TPR-containing Hsc70-Hsp90 organizing protein Hop bridge the activity of Hsp70, which functions on more nascent polypeptide clients, with Hsp90, which folds clients that are closer to their native conformation^32^. Like Hop, DnaJC7 bridges Hsp70 and Hsp90. However, DnaJC7 also enables retrograde transfer of substrates from Hsp90 to Hsp70, allowing additional iterations of the Hsp70-Hsp90 cycle^33^. DnaJC7 thus uniquely facilitates Hsp70/Hsp90- mediated folding and refolding cycles and could be involved in more complex chaperone networks than the canonical JDPs DnaJB1 or DnaJA2.

DnaJC7 complementation studies in the DnaJC7 KO cell lines indicated that expression of WT DnaJC7 in KO cells rescued the tau aggregation phenotype. In contrast, expression of the DnaJC7 (HPQ) mutant, which abolishes DnaJC7:Hsp70 binding^23^, increased seeding relative to all controls, indicating an Hsp70-dependent mechanism for DnaJC7 regulation of tau. By abolishing substrate handoff to Hsp70, the DnaJC7 (HPQ) mutant proteins may become saturated with tau and unable to shuttle tau between Hsp70 and Hsp90, thus disrupting DnaJC7’s role in Hsp70-Hsp90 chaperoning activities and resulting in the apparent dominant negative effects we observed.

### DnaJB6 and DnaJC7 regulation of tau aggregation

JDPs have been implicated in a variety of neurodegenerative diseases. Prior work has suggested that DnaJA2, DnaJB1, and DnaJB4 suppress tau aggregation *in vitro*. The literature has also portrayed JDPs as functionally redundant, with failure of targeted genetic screens knocking out selected JDPs implicated in neurodegeneration to influence tau aggregation^16,34^. We did not identify DnaJA2, DnaJB1, and DnaJB4 as tau aggregation modifiers. This may be because of the model cell line we used.

DnaJB6 oligomers have been reported to potently suppress polyQ, alpha-synuclein, amyloid beta, and TDP-43 aggregation^30,35–38^. Yet DnaJB6 was not previously known to regulate tau aggregation. It remains unknown how DnaJB6 oligomerization impacts its activity, but experiments on DnaJB6 suppression of polyQ indicated that a serine/threonine-rich region may play a role in recognition of substrates^39^. DnaJB6 was not highly abundant in our proteomics screen for tau aggregate interactors, but its KO in tau biosensors increased tau aggregation second only to the DnaJC7 KO. DnaJB6 and its homolog, DnaJB8, have been observed to substoichiometrically inhibit substrate aggregation, suggesting a mechanism based on iterative binding/refolding to aggregation-prone seeds that prevents substrate aggregation^36^.

We have previously reported that DnaJC7 preferentially bound natively folded tau monomer vs. seed-competent monomer or aggregation-prone mutants^15^. In this study, DnaJC7 KO alone increased tau seeding for all seed sources tested, suggesting an interaction with endogenous tau with a mechanism independent of seed conformation. We hypothesize that DnaJC7 suppresses tau aggregation by binding to the inert tau and preventing templating by exogenous tau seeds. This aligns with our previous finding that DnaJC7 preferentially binds to WT tau, which exists in a more closed conformation, than to the P301L mutant tau, which exists in a more open conformation and more closely resembles tau seeds. In conclusion, we identified DnaJC7 as a specific regulator of tau aggregation. It binds tau via its TPR2B domain and engages Hsp70 to stabilize the inert monomer.

## Experimental Procedures

### Identification of aggresome-associated proteins by mass spectrometry

DS1 and DS10 cells were grown to confluency in two T300s per condition. Cells were harvested, pelleted, and washed prior to storage as 4×0.5 T300 pellets at -80°C. For each condition, three pellets were thawed on ice, and each was lysed by trituration in 1 mL ice-cold PBS with 0.25% Triton-X containing cOmplete™ mini EDTA-free protease inhibitor cocktail tablet (Roche) at a concentration of 10% w/vol followed by a 15-minute incubation on ice. Aggresomes and nuclei were collected by centrifuging at 1000xg for 15 minutes followed by resuspension in 400 μL lysis buffer. An Omni-Ruptor 250 probe sonicator was then used at 30% power for thirty 3-second pulses to partially dissolve the pellets. Samples were centrifuged at 250xg for 5 minutes and the supernatant was set aside as Fraction B. Pellets were re-homogenized in an additional 400 μL lysis buffer and sonication and centrifugation was repeated. The final supernatant was added to the previous Fraction B (800 μL volume total). A Bradford assay (Bio-Rad) with BSA standard curve was performed and the protein concentrations were calculated for the nine fractions. Protein concentrations were normalized to 1.1 μg/μL. 72 μL of 10% sarkosyl was added to 650 μL of each sample in ultra-centrifuge tubes (Beckman Coulter) and samples were rotated end-over-end at room temperature for one hour. Samples were then spun at 186,000xg for 60 minutes, supernatant was set aside, and pellets were washed with 1 mL lysis buffer prior to an additional 30 minute 186,000xg spin. Final pellets were resuspended in 30 μL PBS containing 2% SDS and 2% BME by boiling and trituration. 5 μL of Fraction B supernatants and pellets were loaded onto NuPAGE 10% Bis-Tris gels (Life Technologies) and were run at 150 V for 60 minutes. Gels were washed 1x with water and were then stained with SimplyBlue SafeStain (Life Technologies). Images of gels were captured using a digital Syngene imager.

For LC-MS/MS-based detection of proteins, 20 μL re-suspended Fraction B pellets were run 1 cm onto an Any kD Mini-Protean TGX gel (Bio-Rad) followed by Coomassie Blue staining. Whole lanes were excised using ethanol-washed razor blades and gel samples were cut into 1 mm chunks. Gel pieces were reduced with DTT and alkylated with iodoacetamides (Sigma-Aldrich) and were then digested overnight with trypsin (Promega). Next, excised proteins were subjected to solid-phase extraction cleanup with Oasis HLB plates (Waters). The processed samples were then analyzed by LC-MS/MS using a Q Exactive mass spectrometer (Thermo Electron) coupled to an Ultimate 3000 RSLC-Nano liquid chromatography system (Dionex). Samples were injected onto a 180 μm i.d., 15-cm long column packed with reverse-phase material ReproSil-Pur C18-AQ, 1.9 μm resin (Dr. Maisch GmbH, Ammerbuch-Entringen, Germany). Peptides were eluted with a gradient from 1-28% buffer B (80% (v/v) ACN, 10% (v/v) trifluoroethanol, and 0.08% formic acid in water) over 60 minutes. The mass spectrometer could acquire up to 20 MS/MS spectra for each full spectrum obtained. Raw mass spectrometry data files were converted to a peak list format and analyzed using the central proteomics facilities pipeline (CPFP), version 2.0.3^40,41^. Peptide identification was performed using the X!Tandem and open MS search algorithm (OMSSA) search engines against the human protein database from Uniprot, with common contaminants and reversed decoy sequences appended^42,43^. Fragment and precursor tolerances of 20 ppm and 0.1 Da were specified, and three miscleavages were allowed. Carbamidomethylation of Cys was set as a fixed modification and oxidation of Met was set as a variable modification.

Label-free quantitation of proteins across samples was performed using SINQ normalized spectral index software^44^. Finally, spectral counts were added across triplicates. Proteins with a spectral count greater than 5 in DS10, but not identified in DS1 were reported. To calculate enrichment of proteins in the DS10 samples vs DS1 samples, the spectral counts were first negative log_10_ transformed. False Discovery Rate (FDR) analysis was performed using the multiple unpaired t-tests analysis in Prism (GraphPad). The original FDR method of Benjamini and Hochberg was applied, with the desired FDR set to 1.5%. Differences are reported as DS10 spectral index – DS1 spectral index. The -log_10_(q-value) is reported. Gene ontology (GO) analysis was conducted via the Metascape gene annotation and analysis resource^45^.

### Tau Aggregate Degradation Time Courses

OFF1::DS10 cells were treated with two rounds of indicated pooled gRNA lentivirus and were maintained in 24-well plates to assess viability. Two weeks later, a time course examining the decay of tau aggregate seeding activity in the various non-lethal knockouts was performed as follows: Confluent 24-wells were resuspended in 1 mL media and 3.5 μL cells were re-plated into 200 μL total volume in 96-well plates. Tau RD-YFP expression was turned off using 30 ng/mL doxycycline for 1 day, 2 days, 3 days, 4 days, or 5 days. After five days, cells reached confluency and were passaged onto coverslips. 48 hours later, at which point tauRD-YFP expression reached its maximum, cells were fixed and cells containing or lacking inclusions were manually counted. Six replicates of 150+ cells were counted per condition, and averages were calculated. One-way analysis of variance with Bonferroni’s multiple comparison test was used to assess statistical significance relative to non-target controls.

### CRISPR/Cas9 knockout of JDPs in tau biosensor cells

Four human gRNA sequences per gene were selected from the Brunello library^21^. A single non-targeting human gRNA sequence was used as a negative control. For all gRNA sequences not beginning with guanine, a single guanine nucleotide was added at the 5’-end of the sequence to enhance U6 promoter activity. DNA oligonucleotides were synthesized by IDT DNA and cloned into the lentiCRISPRv2 vector^46^ for lentivirus production. The plasmids for the four gRNAs for each gene were pooled together to a final concentration of 40 ng/μL and used to generate lentivirus.

Lentivirus was produced as described previously^47^. Briefly, HEK293T cells were plated at a concentration of 100,000 cells/well in a 24-well plate. 24 hours later, cells were transiently co-transfected with PSP helper plasmid (300 ng), VSV-G (100 ng), and gRNA plasmids (100 ng) using 1.875 μL of TransIT-293 (Mirus) transfection reagent in OptiMEM to a final volume of ∼30 μL. 48 hours later, the conditioned medium was harvested and centrifuged at 1000 rpm for five minutes to remove dead cells and debris. For transduction, 30 μL of the virus suspension was added to HEK293T tau biosensor cells at a cell confluency of 60% in a 96-well plate. 48 hours post-transduction, infected cells were treated with 1 μg/ml puromycin (Life Technologies, Inc.) and maintained under puromycin selection for at least 10 days after the first lentiviral transduction to ensure knockout of the individual JDPs before conducting experiments.

### Recombinant tau seeding assay in JDP knockout tau biosensor cells

The individual JDP knockout cell lines were tested in batches of ten knockout cell lines, including the DnaJC7 and nontargeting control cell lines for each batch. Each batch was assayed in biological duplicate. For all experiments, cells were plated in 96-well plates at 20,000 cells per well in 100 μL of media. 24 hours later, the cells were treated with 50 μL of a heparin-induced recombinant tau fibril dilution series in technical triplicates. Prior to cell treatment, the recombinant tau fibrils were sonicated for 30 seconds at an amplitude of 65 on a Q700 Sonicator (QSonica). A three-fold dilution series of the sonicated fibril concentrations ranging from 100 nM to ∼ 15.2 pM and a media negative control was added to the cell media. 48 hours after treatment with tau, the cells were harvested by 0.05% trypsin digestion and then fixed in PBS with 2% PFA for ten minutes. The cells were then washed and resuspended in 150 μL of PBS.

### Flow cytometry of tau biosensor cells

A BD LSRFortessa was used to perform FRET flow cytometry. To measure mCerulean3 and FRET signal, cells were excited with the 405 nm laser and fluorescence was captured with a 405/50 nm and 525/50 nm filter, respectively. To measure mClover3 signal, cells were excited with a 488 laser and fluorescence was captured with a 525/50 nm filter. To quantify FRET, we used a gating strategy where mCerulean3 bleed- through into the mClover3 and FRET channels was compensated using FlowJo analysis software, as described previously^48^. FRET signal is defined as the percentage of FRET-positive cells in all analyses. For each experiment, 10,000 mClover3/mCerulean3 double-positive cells per replicate were analyzed and each condition was analyzed in triplicate. Data analysis was performed using FlowJo v10 software (Treestar). Two-way analysis of variance with Dunnett’s multiple comparison test was used to assess statistical significance relative to non-target controls in GraphPad Prism.

### Preparation of brain and DS cell lysates

0.5 g frontal lobe sections from AD, PSP, and CBD patients were gently homogenized at 4 °C in 5 mL of TBS buffer containing one tablet of cOmplete™ mini EDTA-free protease inhibitor cocktail (Roche) at a concentration of 20% w/vol using a Dounce homogenizer. Samples were centrifuged at 21,000xg for 15 min at 4ºC to remove cellular debris. Supernatant was partitioned into aliquots, snap frozen in liquid nitrogen, and stored at − 80ºC.

DS1, DS9, and DS10 cell pellets were lysed by resuspension in cold 0.05% Triton in PBS containing cOmplete™ mini EDTA-free protease inhibitor tablet (Roche) at a concentration of 10% w/vol followed by incubation in the lysis buffer for 20 minutes on ice. Homogenates were then clarified by centrifugation at 4ºC at a speed of 500 RCF for 5 minutes followed by 1000 RCF for 5 minutes. The supernatant was then isolated, and total cell lysate protein concentrations were measured using the Pierce™ BCA Protein Assay Kit (ThermoFisher).

### Biosensor seeding with brain and DS cell lysates

Seeding with the brain and DS cell lysates was conducted on the DnaJC7, DnaJB6, and nontargeting knockout tau biosensor cell lines. For all lipofectamine seeding experiments, cells were plated in 96-well plates at 20,000 cells per well in 130 μL of media. 24 hours later, the cells were treated with 20 μL of lipofectamine complexes. The complexes are generated by preparing a lipofectamine in OptiMEM (Gibco) master mix consisting of 1 μL Lipofectamine 2000 (Gibco) and 9 μL OptiMEM per well. The seeding material is prepared in a separate master mix for each lysate tested and consists of 5 μL of brain lysate and 5 μL of OptiMEM per well or 5 μg total protein of DS cell lysate and OptiMEM to 10 μL. The lipofectamine and lysate master mixes are combined in a 1:1 ratio and incubated for 30 minutes at room temperature. The final master mix is then distributed to triplicate wells for the three knockout cell lines, with each well receiving 20 μL of treatment. A lipofectamine control and OptiMEM-only negative control were also generated.

For all naked seeding experiments, cells were plated in 96-well plates at 10,000 cells per well in 100 μL of media. 24 hours later, the cells were treated with 50 μL of seeding complexes. The seeding material is prepared in a separate master mix for each lysate tested and consists of 25 μL of brain lysate and 25 μL of complete media per well or 20 μg total protein of DS cell lysate and media to 50 μL. The final master mix is then distributed to triplicate wells for the three knockout cell lines, with each well receiving 50 μL of treatment. A media-only negative control was also generated.

48 hours after treatment, the cells were harvested by 0.05% trypsin digestion and then fixed in PBS with 2% paraformaldehyde for flow cytometry. Two-way analysis of variance with Dunnett’s multiple comparison test was used to assess statistical significance relative to non-target controls in GraphPad Prism.

### Design of a DnaJC7 construct resistant to targeting by used gRNA sequences

N-terminal Ruby fusion constructs of DnaJC7 were designed to be resistant to targeting by the four gRNAs used to generate the DnaJC7 KO biosensor line. Alternative codon sequences were used at the four gRNA sites to generate a distinct cDNA sequence that maintained the same amino acid sequence. The cDNA sequence is copied below, with alternative codon sequences underlined:

atggcggctgccgcggagtgcgatgtggtaatggcggcgaccgagccggagctgctcgacgaccaagaggcgaagagggaagcagagactttcaaggaacaaggaaatgcatactatgccaagaaagattacaatgaagcttataattattataca aaagccatagatatgtgtcctaaaaatgctagctattatggtaatcgagcagcc<u>acgctgatgatgctgggccgc</u>ttccgg gaagctcttggagatgcacaacagtcagtgaggttggatgacagttttgtccggggacatctacgagag<u>ggtaaatgccat ctgagcctc</u>gggaatgccatggcagcatgtcgcagcttccagagagccctagaactggatcat<u>aagaacgcgcaggcg cagcaggaa</u>ttcaagaatgctaatgcagtcatggaatatgagaaaatagcagaaacagattttgagaagcgagattttcg gaaggttgttttctgcatggaccgtgccctagaatttgcccctgcctgccatcgcttcaaaatcctcaaggcagaatgtttagcaatgctgggtcgttatccagaagcacagtctgtggctagtgacattctacgaatggattccaccaatgcagatgctctgtatgtacgaggtctttgcctttattacgaagattgtattgagaaggcagttcagtttttcgtacaggctctcaggatggctcctgaccacgagaaggcctgcattgcctgcagaaatgccaaagcactcaaagcaaagaaagaagatgggaataaagcatttaagg aaggaaattacaaactagcatatgaactgtacacagaagccctggggatagaccccaacaatataaaaacaaat<u>gcg aagctgtattgcaaccgc</u>ggtacggttaattccaagcttaggaaactagatgatgcaatagaagactgcacaaatgcagt gaagcttgatgacacttacataaaagcctacttgagaagagctcagtgttacatggacacagaacagtatgaagaagcagtacgagactatgaaaaagtataccagacagagaaaacaaaagaacacaaacagctcctaaaaaatgcgcagctggaactgaagaagagtaagaggaaagattactacaagattctaggagtggacaagaatgcctctgaggacgagatcaagaaagcttatcggaaacgggccttgatgcaccatccagatcggcatagtggagccagtgctgaggttcagaaggaggaggagaagaagttcaaggaagttggagaggcctttactatcctctctgatcccaagaaaaagactcgctatgacagtggacaggacctagatgaggagggcatgaatatgggtgattttgatccaaacaatatcttcaaggcattctttggcggtcctggcggcttcagctttgaagcatctggtccagggaatttcttttttcaatttggctga

### Transient overexpression of DnaJC7 constructs in tau biosensor cells

DnaJC7 KO or Non-Targeting control tau biosensor cells were plated at 500K cells/well in a six-well dish. 24 hours later, cells were transiently transfected with 1 ug of plasmid containing the WT, HPQ, or ALS-associated Ruby-tagged sequences of DnaJC7 using 5 μL of Lipofectamine 2000 (Gibco) transfection reagent in OptiMEM (Gibco) to a final volume of 125 μL. The Ruby control cells were transfected with 500 ng of plasmid expressing only Ruby using 5 μL of Lipofectamine 2000 transfection reagent in OptiMEM to a final volume of 125 μL. The vehicle control cells were treated with 5 μL of Lipofectamine 2000 transfection reagent in 120 μL OptiMEM. 48 hours after transfection, the cells were plated in 96-well plates at 20,000 cells per well in 100 μL of media for a standard naked seeding assay. Wells were run on the flow cytometer to completion to ensure collection of a sufficient number of cells with Ruby signal. Two-way analysis of variance with Dunnett’s multiple comparison test was used to assess statistical significance relative to Ruby-alone controls for overexpression and rescue experiments and relative to the WT C7 construct for rescue with ALS-associated mutants in GraphPad Prism.

### Immunoblotting of DnaJC7 from cell lysates

HEK293 tauRD biosensor cell pellets were lysed by resuspension in cold 0.01% Triton in PBS containing cOmplete™ mini EDTA-free protease inhibitor tablet (Roche) at a concentration of 10% w/vol followed by incubation in the lysis buffer for 15 minutes on ice. Homogenates were then clarified by centrifugation at 4ºC at a speed of 17,200 RCF for 15 minutes. The supernatant was then isolated, and total cell lysate protein concentrations were measured using the Pierce™ BCA Protein Assay Kit (ThermoFisher).

10 μg of total cell lysate protein was prepared in 1X (final) LDS Bolt™ buffer (Invitrogen) supplemented with 10% b-mercaptoethanol and heated for 5 minutes at 98ºC. The proteins were resolved by SDS-PAGE using Novex NuPAGE pre-cast gradient Bis-Tris acrylamide gels (4–12%) (Invitrogen). After gel electrophoresis, resolved proteins were transferred onto Immobilon-P PVDF membranes (Millipore Sigma) using a Bio-Rad Trans-blot® semi-dry transfer cell. The membranes were then blocked in 1X TBST buffer (10 mM Tris, 150 mM NaCl, pH 7.4, 0.05% Tween-20) containing 5% non-fat milk powder (Bio-Rad). Membranes were then probed with antibody in TBST containing 5% milk powder.

The following antibodies were used for immunoblotting: rabbit polyclonal anti-DnaJC7 (Proteintech, 11090-1-AP) at a 1:2000 dilution; a secondary donkey-anti-rabbit HRP-linked F(ab’)2 (Cytiva, NA9340-1ML) at a 1:5000 dilution when blotting for DnaJC7; mouse monoclonal anti-Beta-Actin (Proteintech, 66009-1-Ig) at a 1:5000 dilution; and a secondary goat-anti-mouse H&L (HRP) (Abcam, ab6789) at a 1:5000 dilution.

### Calculation of ddG for ALS Mutants

Our structural model of DnaJC7 was built in ROSETTA using homology modeling using DnaJC3 (PDB ID: 3IEG) as a template and minimized using the relax protocol^49,50^. A low scoring model was used to evaluate the energetics of ALS-associated mutations in DnaJC7. The 17 ALS-associated mutations were then individually introduced into the initial DnaJC7 model. For each mutation, 35 replicate simulations were run in parallel for WT and mutant DnaJC7 until convergence. To estimate the ddG of monomer, the mean free energy between the WT and mutant DnaJC7 structures was calculated^29,51^. Simulations were performed on the BioHPC computing cluster at UT Southwestern Medical Center. The relax protocol and Flex ddG used Rosetta v3.13 and v3.12, respectively.

## Acknowledgements

Work in the LAJ lab was supported by an NIH-NIA grant RF1AG078888. Work in the MID lab was supported by the following grants: NIH 3R01AG04678, NIH 1RF1AG059689, NIH1RF1AG065407, The CLW lab was supported by the McCune Foundation and the Winspear Family Center for Research on the Neuropathology of Alzheimer Disease. LAJ, CLW, and MID were supported by the Chan Zuckerberg Initiative 2018-191983, Chan Zuckerberg Initiative 2021-237348. We would like to thank the UT Southwestern Alzheimer’s Disease Center for providing pathological brain tissue samples. We also thank the Proteomics Core Facility and Moody Foundation Flow Cytometry Facility at the University of Texas Southwestern Medical Center.

## Contributions

DWS, VAP, LAJ, and MID conceived and designed the overall study. DWS conducted the proteomics screen and validation of its hits. VAP conducted the genetic screen for all JDPs and the overexpression and rescue experiments with WT and mutant DnaJC7 constructs. AMO generated additional DnaJC7 and DnaJB6 KO cell lines for the time course experiments. BES generated DS cell lysates and aided with brain and cell lysate seeding experiments. VM performed the Rosetta analysis of the ALS-associated mutants. CLW provided patient brain samples. VAP, LAJ, and MID wrote the manuscript.

## Competing Interests

The authors declare that they have no competing interests.

## Data Availability

Raw data for proteomic experiments is unavailable due to a data storage issue. Processed proteomic data and tau aggregate degradation time course data are available on Dryad at doi:10.5061/dryad.fj6q57402. Raw FCS files are deposited on the Cytobank digital repository at https://community.cytobank.org/cytobank/projects/1505.

## Notes

### Competing Interest Statement

The authors have declared no competing interest.

